# M1C is a druggable target for NSCLC KRAS G12C mutant tumors resistant to KRAS inhibitors

**DOI:** 10.64898/2025.12.02.691049

**Authors:** Shinkichi Takamori, Naoki Haratake, Atrayee Bhattacharya, Hiroki Ozawa, Keisuke Shigeta, Mai Onishi, Takefumi Komiya, Hiroki Komatsuda, Tomoyoshi Takenaka, Tomoharu Yoshizumi, Chendi Li, Jiehui Deng, Aaron N. Hata, Kwok K. Wong, Mark D. Long, Donald Kufe

**Author notes:** Corresponding author: Donald Kufe, Dana-Farber Cancer Institute, 450 Brookline Avenue, D830, Boston, MA 02215. Department of Thoracic and Breast Surgery, Oita University Faculty of Medicine, Oita, Japan (Naoki Haratake). University of Texas Health Science Center at Tyler, Tyler, TX, USA.

## Abstract

Treatment of NSCLC KRAS G12C mutant tumors with the allele-selective sotorasib inhibitor is invariably associated with acquired resistance. The MUC1-encoded oncogenic M1C protein is necessary for self-renewal of NSCLC KRAS mutant cells. We report that treatment of NSCLC KRAS G12C cells with sotorasib induces M1C expression by a STAT1-dependent pathway. In turn, M1C drives sotorasib resistance by NF-κB-mediated induction of the epithelial-mesenchymal transition (EMT). Targeting M1C→NF-κB signaling (i) suppresses EMT, and (ii) reverses sotorasib resistance. Of translational relevance, treatment with a M1C antibody-drug conjugate (ADC) is effective against sotorasib-resistant NSCLC KRAS G12C cell line and patient-derived tumor xenografts. Clinically, targeted treatment of patients with NSCLC KRAS G12C tumors overexpressing MUC1 associates with decreases in overall survival. These findings identify M1C as a key effector of sotorasib resistance and as a target for treatment of patients with refractory NSCLC KRAS G12C mutant tumors.

## Introduction

Treatment of NSCLCs harboring the prevalent KRAS G12C mutation has been advanced by development of the sotorasib and adagrasib allele-selective covalent inhibitors^1–3^. In the CodeBreaK 200 phase 3 trial of patients with advanced NSCLC KRAS G12C mutant tumors, sotorasib significantly increased median progression-free survival (PFS) as compared to docetaxel (5-6 months vs 4-5 months)^4^. Moreover, in the KRYSTAL-1 phase 1-2 trial of patients with previously treated NSCLC KRAS G12C tumors, adagrasib achieved a median PFS of 6.5 months and a median overall survival (OS) of 12.6 months^1^. Despite this progress, clinical activity of these agents has been limited by acquired resistance linked to multiple genetic mechanisms that include *KRAS* secondary mutations, *MET* amplification, and activating alterations in *NRAS, BRAF* and *RET*^5–8^. Identification of KRAS switch-II pocket mutations that impair binding of sotorasib and adagrasib uncovered the potential of RAS(ON) tricomplex targeted agents, such as RMC-6236 (NCT05379985) and RMC-7977, to circumvent this resistance^5,6,9–12^. Nonetheless, the pleotropic alterations converging on RAS-MAPK reactivation have limited subsequent treatment options in settings of resistance to KRAS/RAS inhibitors^5–8,10^.

The *MUC1* gene appeared in mammals to protect barrier epithelia lining the respiratory tract from loss of homeostasis^13,14^. *MUC1* is aberrantly expressed in over 80% of NSCLCs and is associated with poor disease-free and overall survival^15,16^. *MUC1* encodes a transmembrane MUC1-C/M1C non-mucin subunit that is activated by inflammation and promotes wound healing^13,14^. Activation of M1C is theoretically reversible with restitution of homeostasis; however, prolonged M1C activation by chronic inflammation induces lineage plasticity and epigenetic reprogramming in association with driving malignant transformation^13,14^. Along these lines, M1C drives EMT, self-renewal capacity and tumorigenicity of NSCLC KRAS mutant cells^17,18^. There is, however, no known involvement of M1C in resistance of NSCLC KRAS G12C mutant cells to KRAS inhibitors.

The present studies show that upregulation of *MUC1* expression in NSCLC KRAS G12C tumors associates with poor overall survival in patients treated with sotograsib and/or adagrasib. Mechanistically, we demonstrate that (i) M1C is induced in the response of NSCLC KRAS G12C cells to these inhibitors, and (ii) M1C confers resistance to KRAS inhibition by activating the NF-κB®ZEB1 pathway and EMT. M1C also activates a mucinous phenotype that has been linked to KRAS G12C inhibitor resistance^5^. Of potential clinical significance, we show that a M1C ADC is effective against sotorasib-resistant NSCLC KRAS G12C cell line and patient-derived tumor xenografts. These findings identify M1C as a target for the treatment of patients with NSCLC KRAS G12C tumors refractory to KRAS inhibition.

## Results

### Targeting NSCLC KRAS G12C signaling induces M1C expression

*MUC1* has been linked to NSCLC progression^15,16^. However, there is no known involvement of the encoded M1C oncogenic subunit in resistance of NSCLC KRAS G12C mutant tumors to KRAS inhibitors. In investigating this unexplored issue, we found that treatment of H358 KRAS G12C cells with 50 and 500 nM of sotorasib for 24 hours results in >5-fold increases in M1C mRNA levels (Fig. 1A). Exposure to 50 nM sotorasib further demonstrated that the induction of M1C transcripts persists through at least day 7 (Fig. 1B). In support of these results, treatment of HCC44 KRAS G12C (Fig. 1C) and H2122 KRAS G12C (Fig. 1D) cells with sotorasib was associated with comparable levels of M1C induction. We also found that treatment of H358 cells with adagrasib increases M1C expression; whereas, interestingly, the KRAS G12D inhibitor MRXT1133 had little if any effect (Fig. 1E). Similar results were obtained with HCC44 (Supplemental Fig. S1A) and H2122 (Supplemental Fig. S1B) cells; that is, treatment with adagrasib, but not MRTX1133, increases M1C transcripts. As a control, treatment of NSCLC H1975 EGFR L858R/T790M cells with the mutant EGFR inhibitor osimertinib induced M1C expression, whereas sotorasib had no apparent effect (Supplemental Fig. S1C). In extending these studies, H358 cells responded to the RAS(ON) inhibitor RMC-7977 with increases in M1C mRNA levels (Supplemental Fig. S1D), indicating that M1C is induced by targeting KRAS, as well as RAS, signaling.

**Figure 1.**
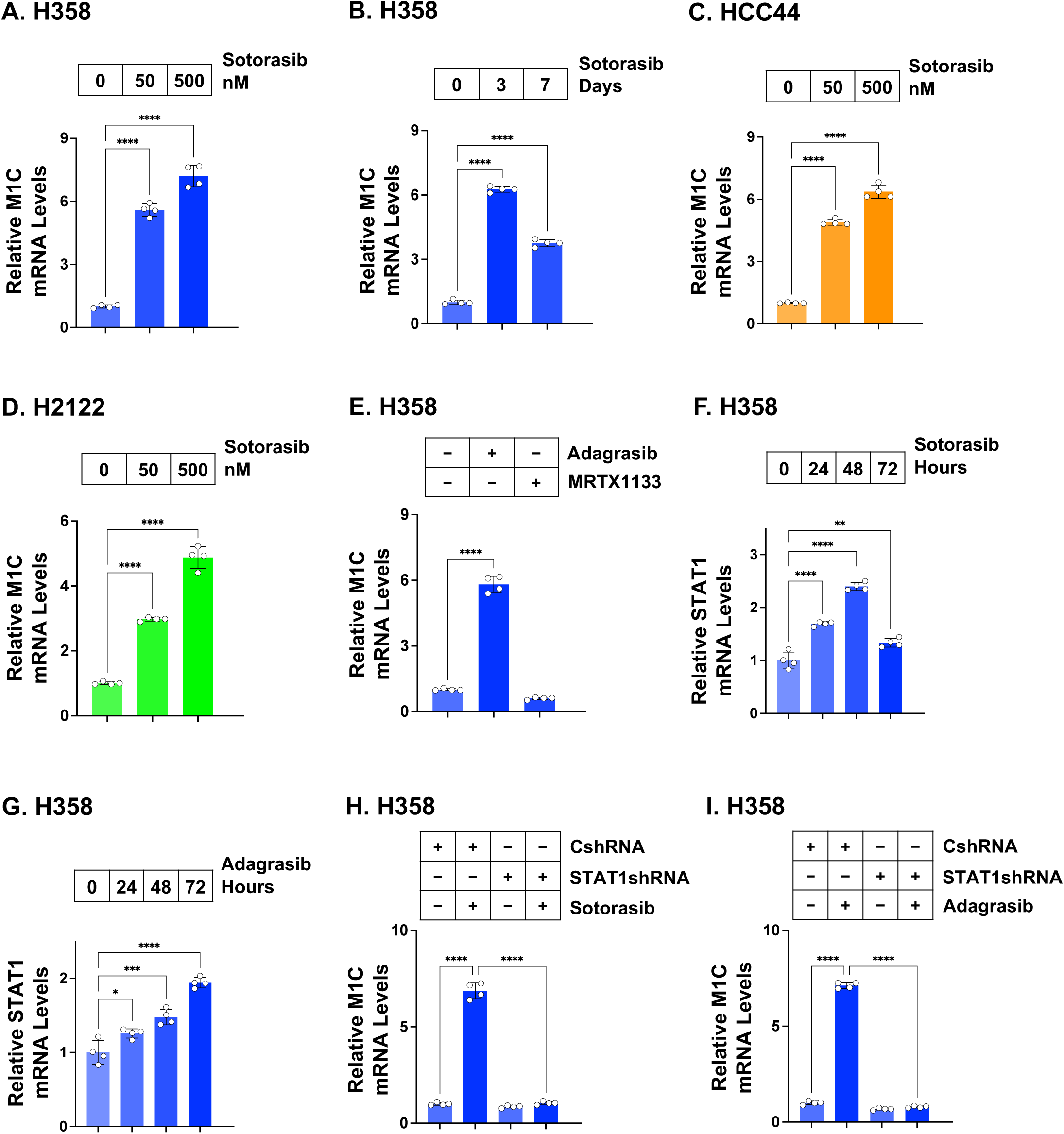
Induction of M1C expression in NSCLC KRAS G12C mutant cells treated with sotorasib and adagrasib. **A.** H358 cells treated with 50 and 500 nM sotorasib for 48 hours were analyzed for M1C transcripts by qRT-PCR using primers listed in Supplemental Table S1. The results (mean±SD of 4 determinations) are expressed as relative levels compared to that obtained for control cells (assigned a value of 1). **B.** H358 cells treated with 50 nM sotorasib for the indicated days were analyzed for M1C transcripts. The results (mean±SD of 4 determinations) are expressed as relative levels compared to that obtained for control cells (assigned a value of 1). **C and D.** HCC44 (**C**) and H2122 (**D**) cells treated with 50 and 500 nM sotorasib for 48 hours were analyzed for M1C transcripts. The results (mean±SD of 4 determinations) are expressed as relative levels compared to that obtained for control cells (assigned a value of 1). **E.** H358 cells treated with 50 nM adagrasib and 50 nM MRTX1133 for 48 hours were analyzed for M1C transcripts. The results (mean±SD of 4 determinations) are expressed as relative levels compared to that obtained for control cells (assigned a value of 1). **F and G.** H358 cells treated with 50 nM sotorasib (**F**) and 50 nM adagrasib (**G**) for the indicated times were analyzed for STAT1 transcripts. The results (mean±SD of 4 determinations) are expressed as relative levels compared to that obtained for control cells (assigned a value of 1). **H and I.** H358/CshRNA and H358/STAT1shRNA cells treated with 50 nM sotorasib (**H**) and 50 nM adagrasib (**I**) for 48 hours were analyzed for M1C transcripts. The results (mean±SD of 4 determinations) are expressed as relative levels compared to that obtained for CshRNA cells (assigned a value of 1).

M1C binds directly to the STAT1 transcription factor (TF) in the response to DNA replication stress and regulates STAT1 target genes, including *MUC1* and *STAT1* in an auto-inductive inflammatory pathway^19,20^. Here, we found that treatment of H358 cells with sotorasib induces STAT1 expression (Fig. 1F). Comparable results were obtained with adagrasib (Fig. 1G), in support of a potential STAT1-dependent mechanism being responsible for induction of M1C expression. Consistent with involvement of this pathway, silencing STAT1 in H358 cells (Supplemental Fig. S1E) abrogated sotorasib (Fig. 1H) and adagrasib (Fig. 1I) -induced M1C expression.

### M1C confers resistance of NSCLC KRAS G12C cells to KRAS inhibition

M1C is expressed as a ∼25 kDa glycoprotein and a 17 kDa unglycosylated protein^13^. Treatment of H358 cells with sotorasib upregulated M1C protein levels (Supplemental Fig. S2A). A similar induction of the M1C protein was observed with RMC-7977 treatment (Supplemental Fig. S2A), indicating that KRAS/RAS inhibition increases M1C mRNA and protein levels. To assess the functional significance of this response, we generated H358 cells expressing a tet-MUC1shRNA and found that DOX treatment suppresses sotorasib-induced (i) M1C expression (Supplemental Fig. S2B), and (ii) killing as assessed by clonogenic survival (Fig. 2A). In further evaluating this M1C dependence, we generated H358 cells stably expressing a second MUC1shRNA#2 and confirmed that downregulating M1C (Supplemental Fig. S2C) suppresses survival in response to sotorasib treatment (Fig. 2B). Analogous findings were obtained in studies with adagrasib (Fig. 2C), indicating that M1C confers resistance to KRAS G12C inhibitors.

**Figure 2.**
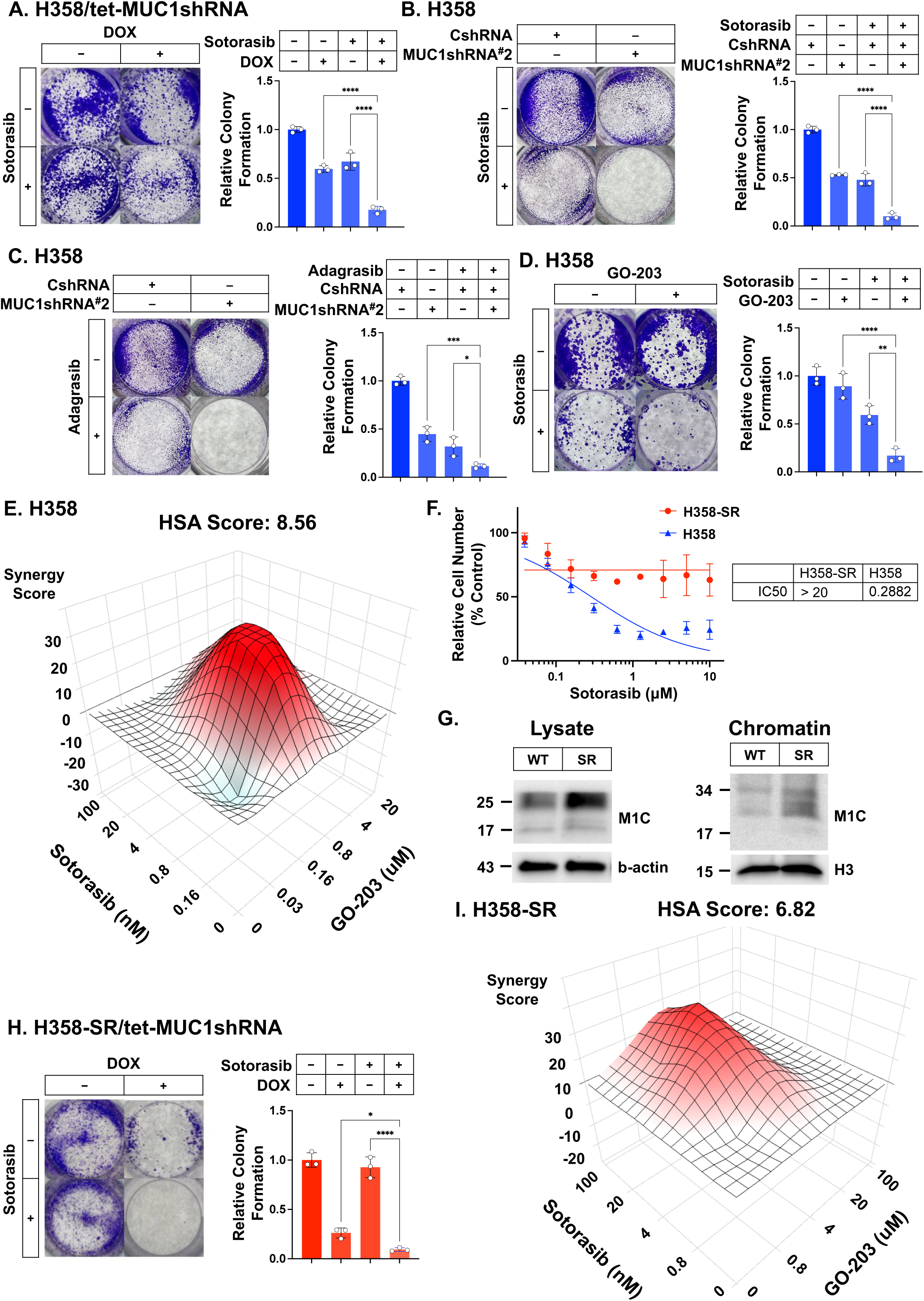
M1C is necessary for sotorasib resistance. **A.** H358/tet-MUC1shRNA cells treated with vehicle of DOX for 7 days and then with 50 nM sotorasib for 7-14 days were analyzed for colony formation. Shown are representative photomicrographs of stained colonies (left). The results (mean±SD of three determinations) are expressed as relative colony formation compared to that for vehicle-treated cells (assigned a value of 1) (right). **B and C.** H358/CshRNA and H358/MUC1shRNA#2 cells treated with 50 nM sotorasib (**B**) and 50 nM adagrasib (**C**) for 7-14 days were analyzed for colony formation. The results (mean±SD of three determinations) are expressed as relative colony formation compared to that for CshRNA cells (assigned a value of 1). **D.** H358 cells treated with 0.5 μM GO-203 and 50 nM sotorasib for 7-14 days were analyzed for colony formation. The results (mean±SD of three determinations) are expressed as relative colony formation compared to that for untreated cells (assigned a value of 1). **E.** H358 cells were treated with the indicated concentrations of GO-203 and sotorasib for 3 days. Cell viability was assessed using Alamar Blue staining (mean of three determinations). Indicated are the combination indices determined using HSA scores. **F.** H358 and H358-SR cells treated with the indicated concentrations of sotorasib for 3 days were assessed for cell viability by Alamar Blue staining (mean of six determinations). Indicated are the IC50 values. **G.** Total cell lysates (left) and chromatin (right) from H358 and H358-SR cells were immunoblotted with antibodies against the indicated proteins. **H.** H358-SR/tet-MUC1shRNA cells treated with vehicle or DOX for 7 days and then with 50 nM sotorasib for 7-14 days were analyzed for colony formation. The results (mean±SD of three determinations) are expressed as relative colony formation compared to that for vehicle-treated cells (assigned a value of 1). **I.** H358-SR cells were treated with the indicated concentrations of GO-203 and sotorasib for 3 days. Cell viability was assessed by Alamar Blue staining (mean of three determinations). Indicated are the combination indices determined using HSA scores.

The M1C cytoplasmic domain contains a CQC motif that is necessary for M1C dimerization and function^21^. Targeting the M1C CQC motif with the GO-203 inhibitor enhanced sensitivity of H358 cells to sotorasib (Fig. 2D) and adagrasib (Supplemental Fig. S2D). We also found that GO-203 synergistically increases sotorasib (Fig. 2E) and adagrasib (Supplemental Fig. S2E) -mediated killing as determined by HSA scores. Whereas these results supported a role for M1C in protecting against KRAS G12C inhibitor-induced loss of survival, we generated sotorasib-resistant H358-SR cells by exposure to increasing sotorasib concentrations over 12 months (Fig. 2F). Of note, H358-SR cells also exhibited resistance to adagrasib (Supplemental Fig. S2F). Analysis of H358-SR cells demonstrated upregulation of M1C levels in total lysates, as well as in chromatin, compared to parental H358 cells (Fig. 2G). Treatment of H358-SR/tet-MUC1shRNA cells with DOX suppressed M1C expression (Supplemental Fig. S2G) and reversed sotorasib resistance (Fig. 2H). In addition, GO-203 synergistically increased sotorasib sensitivity of H358-SR cells (Fig. 2I).

As a second model, we turned to H2122 KRAS G12C mutant cells that are sensitive to sotorasib (IC50=0.7 μM) and generated sotorasib-resistant H2122-SR cells (IC50=16 μM; Supplemental Fig. S2H). Analysis of H2122-SR vs H2122 cells demonstrated upregulation of M1C expression (Supplemental Fig. S2I). Moreover, targeting M1C in H2122-SR cells with GO-203 synergistically increased sotorasib killing (Supplemental Fig. S2J), confirming that M1C is necessary for the sotorasib-resistant phenotype.

### M1C induces EMT and sotorasib resistance

To identify M1C-dependent mechanisms that confer sotorasib resistance, we compared the transcriptomes of parental H358 and H358-SR cells. Silencing M1C in H358 and H358-SR cells downregulated 765 and 206 genes, respectively (Supplemental Figs. S3A-S3C). Among these, we identified 102 genes that included interferon stimulated genes (ISGs), such as IFIT1 and MX1/2, among others (Supplemental Fig. S3C). GSEA of the parental H358 cell RNA-seq dataset uncovered significant M1C-dependent activation of the (i) HALLMARK INTERFERON ALPHA and GAMMA, and (ii) HALLMARK G2M CHECKPOINT, E2F TARGETS and MITOTIC SPINDLE signatures (Fig. 3A), indicative of driving inflammatory and proliferative pathways. Analysis of the H358-SR RNA-seq data further demonstrated that M1C significantly activates the HALLMARK INTERFERON ALPHA and GAMMA, but not the G2M CHECKPOINT, E2F TARGETS and MITOTIC SPINDLE, gene signatures (Fig. 3B). In contrast, we identified M1C-induced activation of the HALLMARK EPITHELIAL MESENCHYMAL TRANSITION signature in H358-SR vs H358 cells (Fig. 3B). Comparison of the H358-SR vs H358 datasets demonstrated marked enrichment of the EMT pathway as compared to other HALLMARK gene signatures (Fig. 3C). Moreover, we confirmed that M1C is necessary for EMT enrichment in H358-SR cells (Fig. 3D; Supplemental Fig. S3D). We also found M1C-dependent enrichment of HALLMARK KRAS SIGNALING UP (Fig. 3E; Supplemental Fig. S3E) and (ii) HALLMARK INFLAMMATORY RESPONSE (Fig. 3F; Supplemental Fig. S3F) gene signatures.

**Figure 3.**
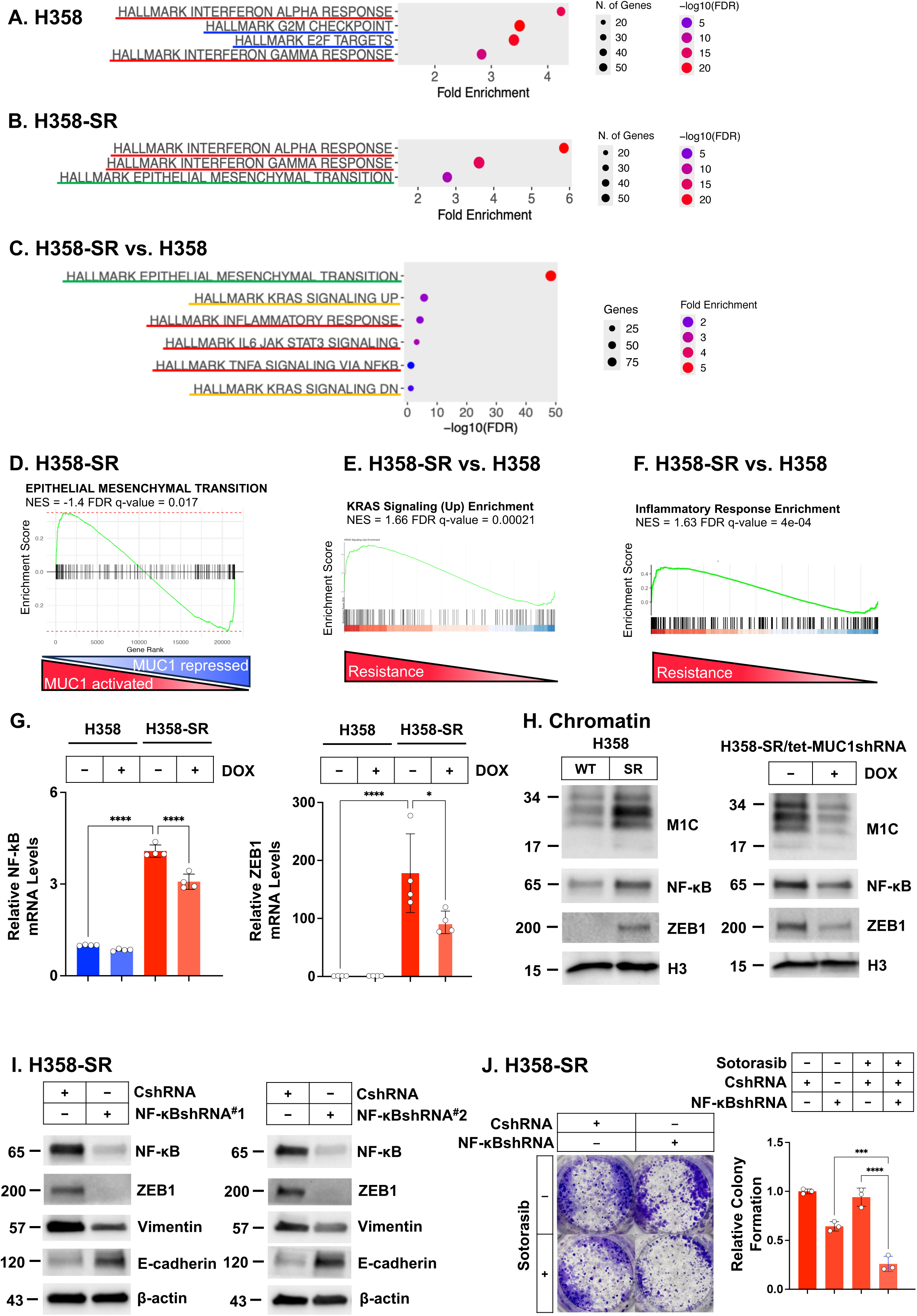
M1C drives sotorasib resistance by a NF-κB/ZEB1/EMT-dependent mechanism. A and. **B.** M1C-dependent activation of the indicated HALLMARK gene signatures in H358 (**A**) and H358-SR (**B**) cells. **C.** Comparison of M1C-induced HALLMARK gene signatures in H358-SR vs H358 cells. **D.** GSEA of H358-SR cells with M1C silencing using the HALLMARK EPITHELIAL MESENCHYMAL TRANSITION gene signature. **E and F.** Comparison of H358-SR vs H358 enrichment using the indicated HALLMARK gene signatures. **G.** H358/tet-MUC1shRNA and H358-SR/tet-MUC1shRNA cells treated with vehicle or DOX for 7 days were analyzed for NF-κB and ZEB1 transcripts. The results (mean±SD of 4 determinations) are expressed as relative levels compared to that obtained for control H358 cells (assigned a value of 1). **H.** Chromatin from H325 and H358-SR cells (left) and H358-SR/tet-MUC1shRNA cells treated with vehicle or DOX for 7 days (right) was immunoblotted with antibodies against the indicated proteins. **I.** Lysates from H358-SR/CshRNA, H358-SR/NF-κBshRNA#1 and H358-SR/NF-κBshRNA#2 cells were immunoblotted with antibodies against the indicated proteins. **J.** H358-SR/CshRNA and H358-SR/NF-κBshRNA cells were treated with 50 nM sotorasib for 7-14 days. Cell viability was assessed by Alamar Blue staining (mean of three determinations).

The ZEB1 EMT TF drives a mesenchymal phenotype in carcinoma cells^22^. M1C binds directly to NF-κB p65 and contributes to the regulation of NF-κB target genes, including ZEB1^23,24^. M1C also binds directly to ZEB1 in driving EMT^24,25^. Analysis of H358-SR vs H358 cells demonstrated M1C-dependent increases in NF-κB p65 and ZEB1 mRNA levels (Fig. 3G). We also found that NF-κB and ZEB1 proteins are increased in chromatin from H358-SR vs H358 cells (Fig. 3H, left) and that silencing M1C downregulates their expression (Fig. 3H, right).

To assess functional involvement of the M1C→NF-κB®ZEB1 pathway, we silenced NF-κB in H358-SR cells with different shRNAs and found (i) downregulation of ZEB1 and (ii) suppression of the EMT phenotype, as evidenced by upregulation of E-cadherin and decreases in vimentin (Fig. 3I). Moreover, as shown for M1C, silencing NF-κB reversed resistance of H358-SR cells to sotorasib (Fig. 3J).

These results were extended with studies of H2122-SR cells, which demonstrated upregulation of M1C, NF-κB and ZEB1 in chromatin, as compared to parental H2122 cells (Supplemental Fig. S3G). In addition, treatment of H2122-SR cells with GO-203 suppressed NF-κB and ZEB1 chromatin levels (Supplemental Fig. S3H). These findings indicate that the M1C→NF-κB®ZEB1 pathway drives EMT in association with sotorasib resistance.

### M1C promotes resistance of patient-derived NSCLC KRAS G12C cells to KRAS inhibition

To extend these results, we studied patient-derived NSCLC KRAS G12C mutant MGH1138 cells, which like H358 cells are sensitive to sotorasib^26^. As found in the H358 and H2122 cell line models, treatment of MGH1138 cells with sotorasib was associated with induction of M1C mRNA (Fig. 4A) and protein (Fig. 4B) levels. Similar results were obtained when treating MGH1138 cells with adagrasib (Supplemental Fig. S4A) and RMC-7977 (Supplemental Fig. S4B). Targeting M1C in MGH1138 cells with GO-203 increased sensitivity to sotorasib as evidenced by significant decreases in clonogenic survival with the combination as compared to that with either agent alone (Fig. 4C). In accordance with these results, GO-203 treatment was synergistic with sotorasib in suppressing MGH1138 cell survival (Fig. 4D).

**Figure 4.**
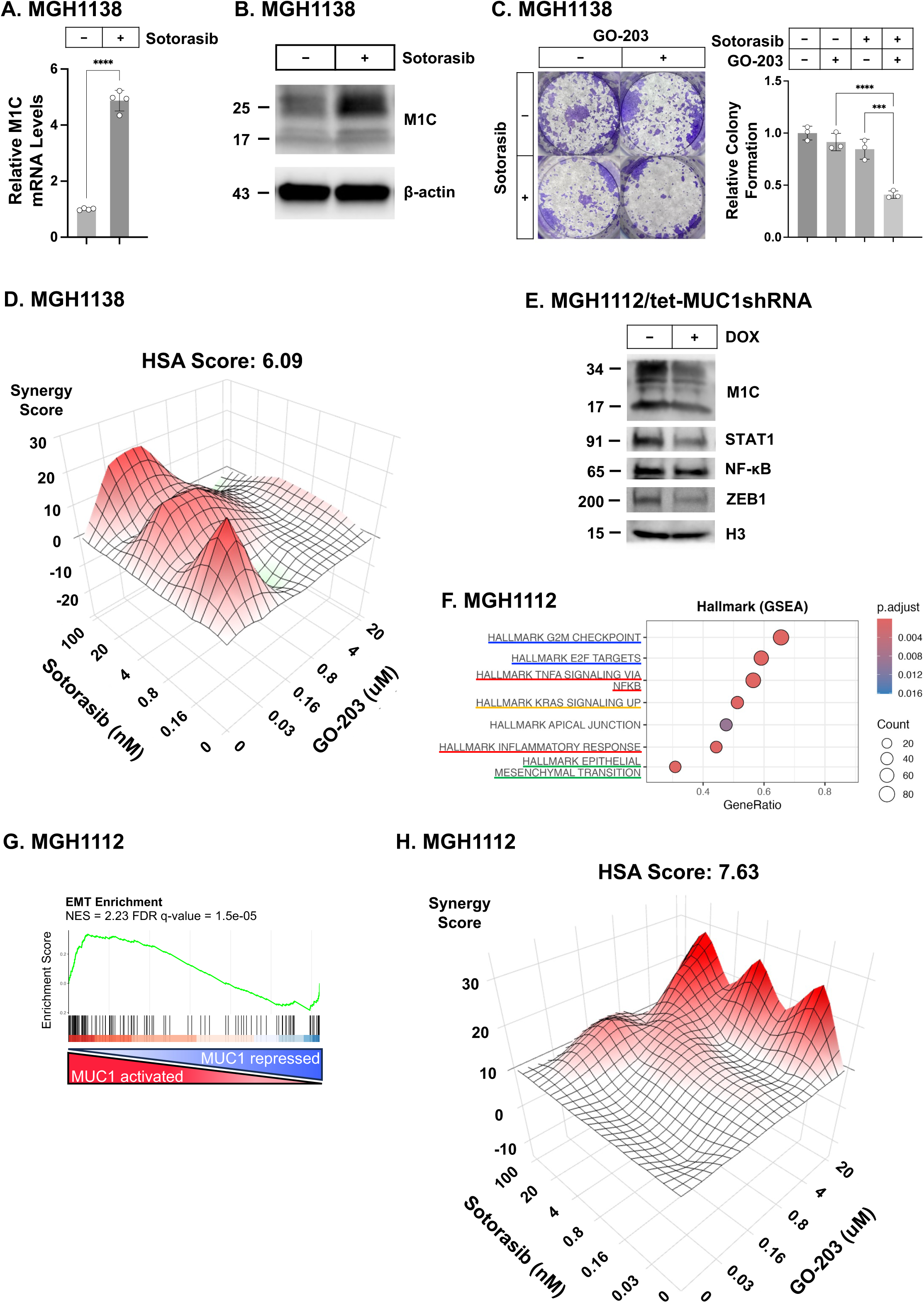
Patient-derived MGH1138 and MGH1112 KRAS G12C cells are M1C-dependent. A and. **B.** MGH1138 cells treated with 50 nM sotorasib for 48 hours were analyzed for M1C transcripts. The results (mean±SD of 4 determinations) are expressed as relative levels compared to that obtained for control cells (assigned a value of 1) **(A).** Lysates were immunoblotted with antibodies against the indicated proteins **(B). C.** MGH1138 cells treated with 0.5 μM GO-203 and 50 nM sotorasib for 7-14 days were analyzed for colony formation. The results (mean±SD of three determinations) are expressed as relative colony formation compared to that for untreated cells (assigned a value of 1). **D.** MGH1138 cells were treated with the indicated concentrations of GO-203 and sotorasib for 3 days. Cell viability was assessed by Alamar Blue staining (mean of three determinations). Indicated are the combination indices determined using HSA scores. **E.** Chromatin from MGH1112/tet-MUC1shRNA cells treated with vehicle or DOX for 7 days was immunoblotted with antibodies against the indicated proteins. **F.** M1C-dependent activation of the indicated HALLMARK gene signatures in MGH1112 cells**. G.** GSEA of RNA-seq data from MGH1112 cells without and with M1C silencing using the HALLMARK EMT gene signature. **H.** MGH1112 cells were treated with the indicated concentrations of GO-203 and sotorasib for 3 days. Cell viability was assessed by Alamar Blue staining (mean of three determinations). Indicated are the combination indices determined using HSA scores.

As a second model, we studied patient-derived NSCLC KRAS G12C MGH1112 cells that, like H358-SR cells, are resistant to sotorasib with an IC50>1 μM^26^. Silencing M1C in MGH1112 cells reduced expression of STAT1, NF-κB and ZEB1 in chromatin (Fig. 4E). GSEA of RNA-seq data from MGH1112 cells revealed that, as shown for H358 cells, silencing M1C suppresses activation of the HALLMARK G2M CHECKPOINT, E2F TARGETS and MITOTIC SPINDLE gene signatures (Fig. 4F). As also found with H358-SR cells, M1C was necessary for activation of the HALLMARK EMT (Fig. 4G), KRAS SIGNALING UP (Supplemental Fig. S4C) and Inflammatory Response (Supplemental Fig. S4D) signatures. Targeting M1C in MGH1112 cells with GO-203 was also synergistic with sotorasib in suppressing survival (Fig. 4H), indicating that M1C is necessary for EMT and the sotorasib-resistant phenotype. In addition, in concert with regulating KRAS signaling, targeting M1C suppressed the feedback p-ERK pathway (Supplemental Fig. S4E).

### Targeting M1C in sotorasib-resistant NSCLC KRAS G12C cell lines with an antibody-drug conjugate

Having found that sotorasib-resistant NSCLC KRAS G12C cells are M1C-dependent, we asked if M1C is a potential target for their treatment. To this end, we assessed the effects of a M1C ADC generated with an antibody against the M1C extracellular domain and conjugated it with a cleavable linker to the MMAE payload^27^. H358, HCC44 and H2122 cells were sensitive to the M1C ADC with IC50 values 84, 62 and 9 nM, respectively (Supplemental Fig. S5A). M1C ADC treatment of H358 cells inhibited clonogenic survival (Fig. 5A) and tumorsphere formation (Fig. 5B). Similar effects were observed with H2122 cells (Supplemental Figs. S5B and S5C), indicating that the M1C ADC is effective in suppressing self-renewal capacity.

**Figure 5.**
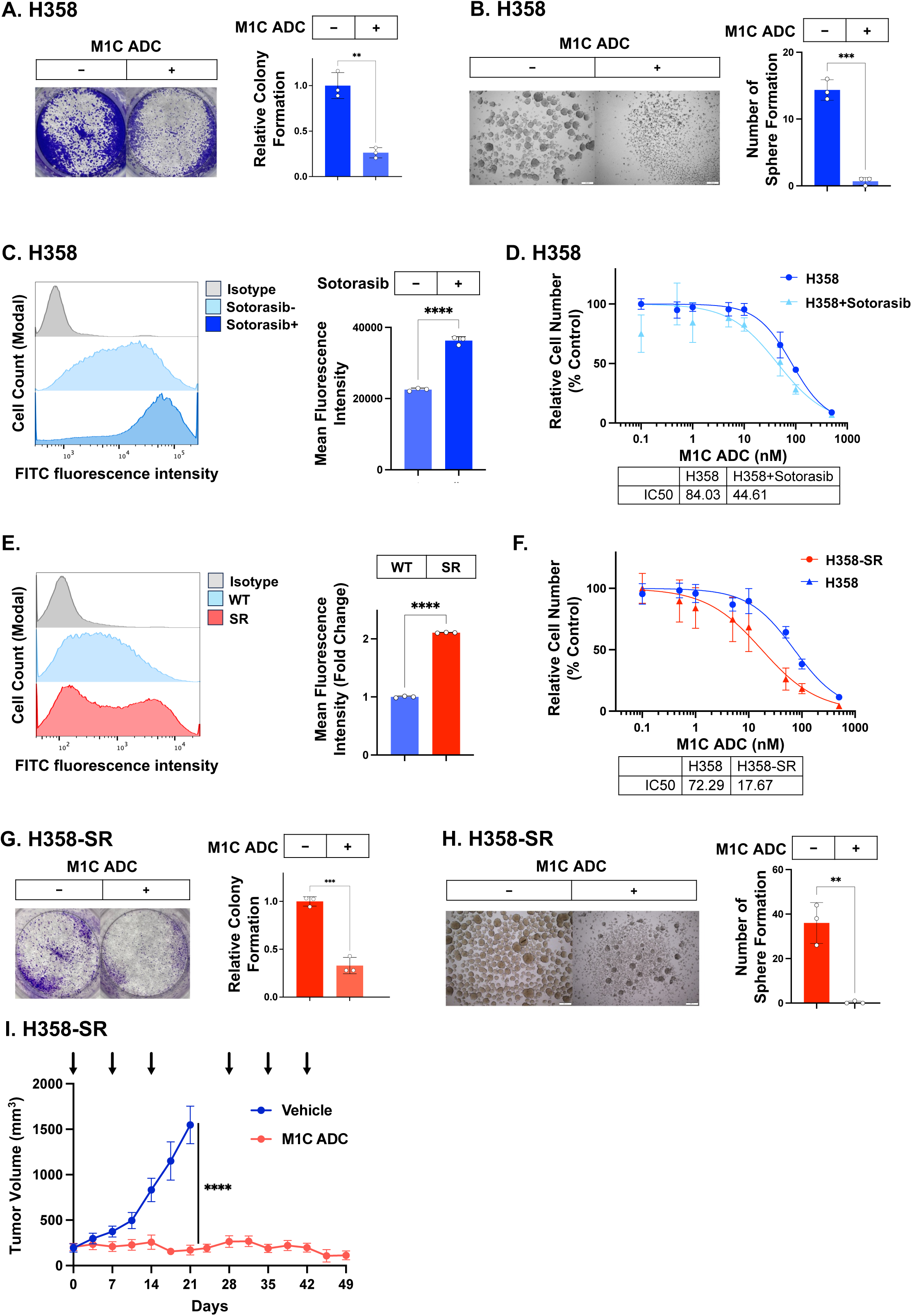
Targeting M1C with an ADC is effective against sotorasib-resistant NSCLC KRAS G12C mutant cells. **A.** H358 cells treated with 20 nM M1C ADC for 7-14 days were analyzed for colony formation. Shown are representative photomicrographs of stained colonies (left). The results (mean±SD of three determinations) are expressed as relative colony formation compared to that for untreated cells (assigned a value of 1) (right). **B.** H358 cells treated with vehicle or 50 nM M1C ADC for 7 days were analyzed for tumorsphere formation. Shown are representative photomicrographs of tumorspheres (left). The results (mean±SD of three determinations) are expressed as number of tumorsphere formations compared to that for vehicle treated cells (right). **C.** H358 cells left untreated or treated with 50 nM sotorasib for 2 days were analyzed by flow cytometry with a control IgG and anti-M1C. Shown are histograms of cells positive for M1C expression (left). The bar plot shows the mean fluorescence intensity (MFI) presented as the mean ± SD of three independent determinations (right). **D.** H358 cells treated with vehicle or 50 nM sotorasib for 2 days and then M1C ADC for 7 days were analyzed for cell viability by Alamar blue staining. **E.** H358 and H358-SR cells were analyzed by flow cytometry with a control IgG and anti-M1C. Shown are histograms of cells positive for M1C expression (left). The bar plot shows the MFI, presented as the mean ± SD of three independent determinations (right). **F.** H358 and H358-SR cells treated with the indicated concentrations of M1C ADC for 7 days were analyzed for cell viability by Alamar Blue staining. The results (mean±SD of six determinations) are expressed as relative cell number (% control) compared to that for untreated cells. Indicated are the M1C ADC IC50 values. **G.** H358-SR cells treated with 50 nM M1C ADC for 7-14 days were analyzed for colony formation. Shown are representative photomicrographs of stained colonies (left). The results (mean±SD of three determinations) are expressed as relative colony formation compared to that for untreated cells (assigned a value of 1) (right). **H.** H358-SR cells treated with vehicle or 50 nM M1C ADC for 7 days were analyzed for tumorsphere formation. Shown are representative photomicrographs of tumorspheres (left). The results (mean±SD of three determinations) are expressed as number of tumorsphere formations compared to that for vehicle treated cells (assigned a value of 1) (right). **I.** Nude mice were injected subcutaneously with 5 × 10^6^ H358-SR cells. Mice were randomized into two groups when the mean tumor volume reached 100 mm^3^ and then treated with vehicle (n = 6) or 7.5 mg/kg M1C ADC weekly x 3 every 28 days for 2 cycles (n = 6). Tumor volumes are expressed as the mean±SEM.

Consistent with the demonstration that sotorasib induces M1C expression, we found that sotorasib and adagrasib increase M1C levels on the surface of H358 cells (Fig. 5C; Supplemental Fig. S5D) and sensitivity to the M1C ADC (Figs. 5D and Supplemental Fig. S5E). Interestingly, we also found that (i) M1C expression is significantly increased on the surface of H358-SR vs H358 cells (Fig. 5E), and (ii) H358-SR cells are more sensitive to M1C ADC treatment (IC50=18 nM) than H358 cells (IC50=72 nM)(Fig. 5F). The M1C ADC was also more effective against H2122-SR vs H2122 cells (Supplemental Fig. S5F), indicating that sotorasib resistance is associated with increased sensitivity to this agent.

These results were extended by the demonstration that the M1C ADC suppresses H358-SR (Fig. 5G) and H2122-SR (Supplemental Fig. S5G) clonogenic survival. The M1C ADC was also highly effective in inhibiting H358-SR (Fig. 5H) and H2122-SR (Supplemental Fig. S5H) tumorsphere formation. Given these results, we treated established H358-SR tumor xenografts with the M1C ADC and found induction of complete and durable responses (Fig. 5I) in the absence of weight loss or other overt toxicities (Supplemental Fig. S5I).

### M1C is a potential target for ADC treatment of patients with NSCLC KRAS G12C tumors resistant to KRAS inhibitors

In extending these results with patient-derived cells, we found that M1C ADC treatment of MGH1138 cells is effective in suppressing clonogenic survival (Fig. 6A). Additionally, treatment of MGH1138 cells with the M1C ADC in combination with sotorasib demonstrated synergistic activity (Fig. 6B). We also found that exposure of MGH1138 cells to the M1C ADC is effective in inhibiting tumorsphere formation (Fig. 6C), indicating that the ADC could be used alone and in combination with sotorasib to suppress self-renewal capacity.

**Figure 6.**
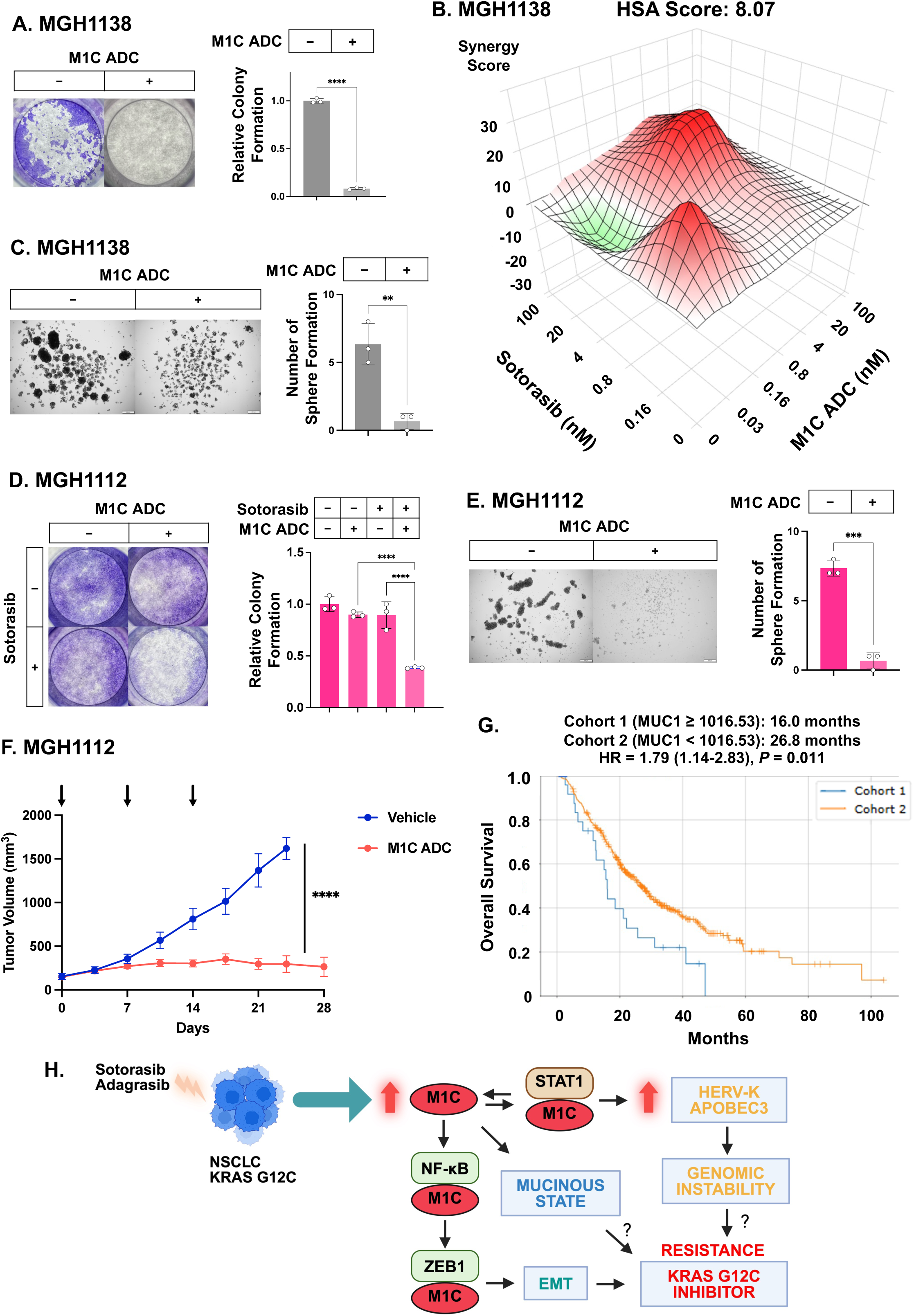
M1C is a potential target for the treatment of human NSCLC KRAS G12C tumors with the M1C ADC. **A.** MGH1138 cells treated with 50 nM M1C ADC for 7 days were analyzed for colony formation. Shown are representative photomicrographs of stained colonies (left). The results (mean±SD of three determinations) are expressed as relative colony formation compared to that for untreated cells (assigned a value of 1) (right). **B.** MGH1138 cells were treated with the indicated concentrations of M1C ADC and sotorasib for 7 days. Cell viability was assessed by Alamar Blue staining (mean of three determinations). Indicated are the combination indices determined using HSA scores. **C.** MGH1138 cells treated with vehicle or 50 nM M1C ADC for 7 days were analyzed for tumorsphere formation. Shown are representative photomicrographs of tumorspheres (left). The results (mean±SD of three determinations) are expressed as number of tumorsphere formations compared to that for vehicle treated cells (right). **D.** MGH1112 cells treated with vehicle or 0.5 µM sotorasib and 5 nM M1C ADC for 7-14 days were analyzed for colony formation. Shown are representative photomicrographs of stained colonies (left). The results (mean±SD of three determinations) are expressed as relative colony formation compared to that for untreated cells (assigned a value of 1) (right). **E.** MGH1112 cells treated with vehicle or 50 nM M1C ADC for 7 days were analyzed for tumorsphere formation. Shown are representative photomicrographs of tumorspheres (left). The results (mean±SD of three determinations) are expressed as number of tumorsphere formations compared to that for vehicle treated cells (right). **F.** Nude mice were injected subcutaneously with 5 × 10^6^ MGH1112 cells. Mice were randomized into two groups when the mean tumor volume reached 100 mm^3^ and then treated with vehicle (n = 6) or 7.5 mg/kg M1C ADC weekly x 3 every 28 days for 2 cycles (n = 6). Tumor volumes are expressed as the mean±SEM. **G.** Overall survival for patients with NSCLC G12C mutant tumors treated with sotorasib and/or adagrasib according to MUC1 high (cohort 1) and MUC1 low (cohort 2) levels using the CARIS CODEai database. **H.** Schema depicting involvement of M1C in conferring resistance of NSCLC KRAS G12C cells to KRAS inhibition. Treatment of NSCLC KRAS G12C cells with sotorasib or adagrasib induces M1C expression by activation of the M1C/STAT1 auto-inductive pathway. M1C/STAT1 signaling activates HERV and APOBEC3 expression, which drives genomic instability and drug resistance^33^. M1C is also induced by the RAS(ON) inhibitor RMC-7977, but not MRTX1133, indicating that this induction represents a response to both KRAS/RAS inhibition. Upregulation of M1C activates NF-κB®ZEB1 signaling, which involves direct interactions of the M1C cytoplasmic domain with NF-κB and ZEB1. In this way, M1C integrates the inflammatory STAT1/mucinous state with the NF-κB®ZEB1®EMT pathway. These results support a model in which M1C confers resistance to KRAS G12C inhibition by epigenetically integrating chronic inflammation and the induction of EMT.

In studies of sotorasib-resistant MGH1112 cells, the M1C ADC was active in suppressing clonogenic survival (Supplemental Fig. S6A). Moreover, the M1C ADC was more effective in combination with sotorasib than either agent alone (Fig. 6D; Supplemental Fig. S6B). Importantly, the M1C ADC was also highly effective in inhibiting MGH1112 self-renewal capacity as evidenced by suppression of tumorsphere formation (Fig. 6E). In addition, treatment of established MGH1112 tumor xenografts with the M1C ADC induced durable regressions (Fig. 6F) in the absence of significant weight loss (Supplemental Fig. S6C).

Based on these results, we examined M1C dependence in driving a mucinous transcriptional state that associates with survival of NSCLC KRAS G12C tumors treated with RAS pathway inhibitors^5^. The mucinous state includes expression of (i) secreted and transmembrane mucins that form a protective physical barrier and (ii) trefoil secreted peptides, such as TFF1, that contribute to the mucous layer in protecting against biotic and abiotic insults^5,28,29^. Silencing M1C in H358 cells decreased expression of the secreted MUC3 mucin, the transmembrane MUC4, MUC13, MUC16 and MUC20 mucins, and TFF1 (Supplemental Fig. S6D). M1C was also necessary for expression of mucin genes and TFF1 in MGH1112 cells (Supplemental Fig. S6E). By extension, analysis of the KRYSTAL-1 trial dataset demonstrated that MUC1 associates with mucin gene and TFF1 expression, indicating that M1C drives the mucinous, as well as mesenchymal, states. In further investigating the potential of M1C as a therapeutic target, we analyzed the CARIS CODEai database and found that, when treated with sotorasib and/or adagrasib, patients with NSCLC KRAS G12C tumors expressing high vs low MUC1 levels exhibit a significant decrease in overall survival (Fig. 6G).

## Discussion

Acquired resistance to sotorasib and adagrasib has been linked to multiple genetic alterations^5–8,10,30^. The present results uncover a non-genetic mechanism in which M1C is activated in the response of NSCLC KRAS G12C mutant cells to treatment with these inhibitors (Fig. 6H). M1C interacts with RTKs at the NSCLC cell membrane and functions as a scaffold for transducing downstream signals^13^. In this way, the M1C cytoplasmic domain forms complexes with SHP2, PI3K, SHC, PLCg, and GRB2/SOS that integrate RTK and RAS signaling pathways^31^. Interestingly, this induction of M1C is not restricted to inhibiting KRAS G12C, as it extends to inhibiting RAS more broadly with the RAS(ON) RMC-7977 inhibitor. Studies in PDAC cells have further demonstrated that M1C is induced by KRAS G12D MRTX1133 inhibitor^32^, indicating that M1C activation plays a non-genetic role in compensating to KRAS/RAS inhibition. The induction of M1C in response to KRAS inhibition is consistent with the evolution of this protein in mammals to protect barrier tissues from loss of homeostasis^13,31^. M1C induces inflammatory pathways that drive wound repair and are reversible with resolution of the insult^13,31^. As a result, the *MUC1* gene is activated by an inflammatory memory response to DNA replication stress induced by targeted agents^33,34^. Conversely, as an adverse adaptation, prolonged M1C activation by chronic inflammation promotes malignant progression^13,31^. Our findings demonstrate that inhibition of KRAS G12C signaling induces M1C by a STAT1-dependent pathway (Fig. 6H), which has been linked to activation of (i) IFN type I/II signaling, (ii) HERV, APOBEC3 and other ISG expression, (iii) inflammatory memory, and (iv) drug resistance^19,33–36^.

The finding that M1C is activated by KRAS G12C inhibitors invoked the possibility that upregulation of M1C may be responsible for the drug-resistant phenotype. The STAT and NF-κB pathways intersect in complex regulatory networks in association with driving chronic inflammation in cancer^37^. In KRAS G12C inhibitor-resistant cell models, we found that M1C, as well as NF-κB and ZEB1, levels are upregulated in chromatin. Like STAT1, M1C forms a direct complex with NF-κB and regulates NF-κB target genes^33^. As a result, M1C/NF-κB complexes induce *ZEB1* transcription and, in turn, M1C directly activates ZEB1^24^. Here, we found that the M1C/NF-κB pathway is responsible for progression of wild-type NSCLC KRAS G12C mutant cells to the KRAS inhibitor-resistant phenotype by driving ZEB1 and the EMT state (Fig. 6H). These findings support a model in which (i) M1C is induced by STAT1 in sotorasib-treated parental cells, and (ii) M1C→NF-κB®ZEB1 signaling drives EMT and resistance to KRAS G12C inhibition (Fig. 6H). The M1C cytoplasmic domain activates JAK1-STAT1^20,38^, TAK1-NF-κB^23,39^ and ZEB1^24^ signaling through the same binding regions, indicating that their induction is conferred by integrated M1C-dependent inflammatory mechanisms. Our results further demonstrate that, in addition to driving inflammation and EMT, M1C is necessary for induction of a mucinous state characterized by expression of (i) secreted and transmembrane mucins^29^, and (ii) TFF1^28^. Expression of genes in the mucinous/epithelial and mesenchymal states has been linked to resistance of NSCLC KRAS G12C tumors to adagrasib^5^. The functional significance of these distinct states along the epithelial-mesenchymal axis is unclear^5^, but could relate to activation of M1C signaling in promoting the partial-EMT (p-EMT) phenotype^40^ (Fig. 6H).

Patients with NSCLC KRAS G12C tumors resistant to sotorasib and/or adagrasib have few treatment options^6–8,10^. Having identified that M1C is a potential target for the treatment of cells resistant to KRAS G12C inhibitors, we asked if a M1C ADC is effective in this setting. Our results demonstrate that the M1C ADC is active in vitro against NSCLC KRAS G12C parental, as well as sotorasib-resistant, cells. In addition, the M1C ADC was highly effective against H358-SR and patient-derived sotorasib-resistant MGH1112 xenograft tumors. Of further relevance as a potential target, we found that upregulation of MUC1 expression in NSCLC KRAS G12C tumors associates with decreases in overall survival for patients treated with sotorasib/adagrasib. These findings collectively lend support for development of the M1C ADC for treatment of sotorasib/adagrasib-resistant tumors. The M1C ADC has also been shown to be effective drug-resistant (i) HR+/HER2- breast^41^ (ii) CRPC/NEPC prostate^42^ and (iii) PDAC KRAS G12D^43^ tumor models, indicating that this agent may be broadly active in settings of tumors unresponsive to targeted agents. Along these lines, the M1C ADC is under development by the NCI NExT Program for IND-enabling studies with the plan for the NCI CTEP Program to evaluate this agent in early phase clinical trials. In summary, the present results demonstrate that M1C is necessary for KRAS G12C inhibitor resistance and is a potential target for treatment of patients with refractory NSCLC KRAS G12C tumors.

## Materials and Methods

### Cell culture

H358 KRAS G12C mutant, TP53 null (ATCC, Manassas, VA, USA), HCC44 KRAS G12C mutant, TP53 null (ATCC) and H2122 KRAS G12C mutant, TP53 null (ATCC) cells were cultured in RPMI 1640 medium (Corning, NY, USA) supplemented with 10% FBS and 2 mM glutamine. MGH1138 (KRAS G12C, STK11 mutant) and MGH1112 (KRAS G12C, STK11 and KEAP1 mutant) cells obtained from Aaron Hata were maintained in RPMI1640 medium with 10% FBS as described^26^. Sotorasib-resistant cells maintained in the presence of drug were grown in drug-free medium for 24 h before analysis. Authentication of the cells was performed by short tandem repeat (STR) analysis every 4 months as described^19^. Cells were monitored for mycoplasma contamination using the MycoAlert Mycoplasma Detection Kit (Lonza, Rockland, ME, USA) every 3 months.

### Gene silencing

MUC1shRNA (MISSION shRNA TRCN0000122938; Sigma) or a control scrambled shRNA (CshRNA; Sigma) was inserted into the pLKO-tet-puro vector (Plasmid #21915; Addgene, Cambridge, MA, USA). MUC1shRNA#2 (MISSION shRNA TRCN0000430218), STAT1shRNA (TRCN0000004266) and, NF-κBshRNA#1 (TRCN000014686), and NF-κBshRNA#2 (TRCN000014687) were produced in HEK293T cells as described^19^. Cells transduced with the vectors were selected for growth in 1-4 mg/ml puromycin as described^19^. Cells were treated with 0.1% DMSO as the vehicle control or 500 ng/ml doxycycline (DOX; Millipore Sigma).

### Quantitative reverse-transcription PCR (qRT-PCR)

Total RNA was extracted using TRIzol reagent (Thermo Fisher Scientific, Waltham, MA, USA) as described^19^. cDNA synthesis was performed with the High-Capacity RNA-to-cDNA Kit (Applied Biosystems, Grand Island, NY, USA). Quantitative PCR was performed using Power SYBR Green PCR Master Mix (Applied Biosystems) as described^19^. Primer sequences are listed in Supplementary Table S1.

### Immunoblot analysis

Total cell lysates and chromatin prepared from non-confluent cells were analyzed by immunoblotting with anti-M1C (16564, 1:1000 dilution; Cell Signaling Technology (CST), Danvers, MA, USA, RRID:AB_2798765), anti-β-actin (A5441, 1:2000 dilution; Sigma-Aldrich, Burlington, MA, USA, RRID:AB_476744), anti-Histone H3 (9715, 1:1000 dilution; CST, RRID:AB_331563), anti-STAT1 (RAB01893, 1:500 dilution; CST, RRID:AB_3710444), anti-ERK (9107, 1:1000 dilution; CST, RRID:AB_10695739), anti-p-ERK (4377, 1:1000 dilution; CST, RRID:AB_331775), anti-NF-κB p65 (8242, 1:1000 dilution; CST, RRID:AB_10859369), anti-ZEB1 (3396, 1:1000 dilution; CST, RRID:AB_1904164), anti-E-cadherin (3195, 1:1000 dilution; CST, RRID:AB_2291471) and anti-vimentin (5741, 1:1000 dilution; CST, RRID: AB_10695459) as described^19^.

### RNA-seq analysis

Total RNA from cells cultured separately in triplicates was isolated using Trizol reagent (Invitrogen) as described^19^.

### Cell viability analysis

One to 1-2 thousand cells were seeded per well in 96-well plates (Thermo Fisher Scientific, Waltham, MA, USA) and incubated for 24 hours before treatment. Cell viability was assessed using Alamar Blue staining (Thermo Fisher Scientific). Synergistic effects were evaluated using the HSA model implemented in SynergyFinder Plus 3.0. The HSA independence model was used to determine if the drug combination effect is equal to, higher than, or lower than the expected effect (E_AB_) and thereby the combination is deemed additive (HSA = 0), synergistic (> 0), or antagonistic (< 0), respectively.

### Clonogenic survival assays

Cells were seeded at 2000 cells/well in 24-well plates and treated after 24 hours of culture. The cells were stained with 0.5% crystal violet in 25% methanol on day 7-14 after treatment and evaluated for colony formation as described^19^.

### Tumorsphere formation assays

Cells (5 x 10^3^) were seeded per well in 6-well ultra-low attachment culture plates (Corning Life Sciences, Corning, NY, USA) and cultured as described^19^. Tumorspheres with a diameter >200 μm were counted under an inverted microscope in triplicate wells.

### Flow cytometry

Cells were harvested and incubated with either mAb 3D1 or an IgG1 kappa isotype control antibody (cat# 60070.1; STEMCELL Technologies, Vancouver, BC, Canada). Goat anti-mouse IgG (Alexa Fluor 488) (Abcam, Cambridge, MA, USA, ab150113; 1:100 dilution) was used as the secondary antibody. Data were acquired and analyzed as described^43^.

### Sex as a biological variable

Our animal study exclusively examined female mice. It is unknown whether the findings are relevant for male mice.

### Mouse tumor model studies

Six-week-old nude female mice (The Jackson Laboratory; Bar Harbor, ME, USA) were injected subcutaneously in the flank with 2-5 x 10^6^ H358-SR or MGH1112 cells in 100 μl of a 1:1 solution of medium and Matrigel (BD Biosciences). Mice were pair-matched into treatment groups when the mean tumor volumes reached 100-150 mm^3^. Tumor measurements and body weights were recorded every 3-4 days. Mice were sacrificed when tumors reached >1000 mm^3^ as calculated by the formula: (width)^2^ x length/2. These studies were conducted in accordance with ethical regulations required for approval by the Dana-Farber Cancer Institute Animal Care and Use Committee (IACUC) under protocol 03-029.

### Statistical analysis

Each experiment was performed at least three times. Data are expressed as the mean±SD. The unpaired Mann-Whitney U test was used to determine differences between means of groups. Asterisks represent *P ≤ 0.05, **P ≤ 0.01, ***P ≤ 0.001, ****P ≤ 0.0001 with CI = 95%.

## Data Availability

The RNA-seq datasets have been submitted to the GEO for GSE numbers once government funding is restored. Additional raw data is available from the corresponding author upon reasonable request.

## Acknowledgements

Research reported in this publication was supported by the National Cancer Institute of the National Institutes of Health under grant numbers CA97098, CA282437, CA289134 awarded to DK. This project has also been supported through the National Cancer Institute Experimental Therapeutics Program (NExT).

## Author contributions

Conceptualization: ST, NH, AB, HO, KS, MM; Methodology: ST, NH, AB, HO, KS, MM, HK, CL, JD; Investigation: ST, AB, HO, KS, MM, TK, ML; Writing – original draft: DK; Writing – review & editing: ST, NH, AB, TT, TY, AH, KW, ML, DK; Supervision: AH, KW, DK; Funding acquisition: DK.

## Author Disclosures

DK has equity interests in Genus Oncology.

## Additional information

**Correspondence** and requests for materials should be addressed to: Donald Kufe, Dana-Farber Cancer Institute, 450 Brookline Avenue, D830, Boston, MA 02215.

---

**Supplemental Figure S1.** Targeting KRAS induces M1C expression. A and. **B.** HCC44 (**A**) and H2122 (**B**) cells treated with 50 nM adagrasib and 50 nM MRTX1133 for 48 hours were analyzed for M1C transcripts. The results (mean±SD of 4 determinations) are expressed as relative levels compared to that obtained for control cells (assigned a value of 1). **C.** H1975 EGFR mutant cells treated with 50 nM osimertinib and 50 nM sotorasib for 48 hours were analyzed for M1C transcripts. The results (mean±SD of 4 determinations) are expressed as relative levels compared to that obtained for control cells (assigned a value of 1). **D.** H358 cells treated with 100 nM RMC-7977 for the indicated times were analyzed for M1C transcripts. The results (mean±SD of 4 determinations) are expressed as relative levels compared to that obtained for control cells (assigned a value of 1). **E.** H358/CshRNA and H358/STAT1shRNA cells were analyzed for STAT1 transcripts. The results (mean±SD of 4 determinations) are expressed as relative levels compared to that obtained for CshRNA cells (assigned a value of 1).

**Supplemental Figure S2.** Targeting M1C reverses sotorasib resistance. **A.** Lysates from H358 cells treated with 50 nM sotorasib for 48 hours and 100 nM RMC-7977 for 48 hours were immunoblotted with antibodies against the indicated proteins. **B.** Lysates from H358/tet-MUC1shRNA cells treated with vehicle or DOX for 7 days and then 50 nM sotorasib for 48 hours were immunoblotted with antibodies against the indicated proteins. **C.** Lysates from H358/CshRNA and H358/MUC1shRNA#2 cells treated with vehicle or 50 nM sotorasib for 48 hours were immunoblotted with antibodies against the indicated proteins. **D.** H358 cells treated with 0.5 μM GO-203 and 50 nM adagrasib for 7-14 days were analyzed for colony formation. The results (mean±SD of three determinations) are expressed as relative colony formation compared to that for untreated cells (assigned a value of 1). **E.** H358 cells were treated with the indicated concentrations of GO-203 and adagrasib for 3 days. Cell viability was assessed by Alamar Blue staining (mean of three determinations). Indicated are the combination indices as determined using HSA scores. **F.** H358 and H358-SR cells treated with the indicated concentrations of adagrasib for 3 days were assessed for cell viability by Alamar Blue staining (mean of six determinations). Indicated are the IC50 values. **G.** Lysates from H358-SR/tet-MUC1shRNA cells treated with vehicle or DOX for 7 days were immunoblotted with antibodies against the indicated proteins. **H.** H2122 and H2122-SR cells treated with the indicated concentrations of sotorasib for 3 days were assessed for cell viability by Alamar Blue staining (mean of six determinations). Indicated are the IC50 values. **I.** Lysates from H2122 and H2122-SR cells were immunoblotted with antibodies against the indicated proteins. **J.** H2122-SR cells were treated with the indicated concentrations of GO-203 and sotorasib for 3 days. Cell viability was assessed by Alamar Blue staining (mean of three determinations). Indicated are the combination indices determined using HSA scores.

**Supplemental Figure S3.** M1C regulates transcriptomes of H358 and H358-SR cells. A and. **B.** Volcano plots of down- and upregulated genes in H358 (**A**) and H358-SR (**B**) cells with M1C silencing. **C.** Venn diagram of down- and upregulated genes in H358 and H358-SR cells with M1C silencing. **D.** Comparison of H358-SR vs H358 enrichment using the HALLMARK EMT gene signature. **E and F.** GSEA of H358-SR cells with M1C silencing using the indicated HALLMARK gene signatures. **G.** Chromatin from H2122 and H2122-SR cells was immunoblotted with antibodies against the indicated proteins. **H.** Lysates from H2122-SR cells treated with 5 μM GO-203 for 48 hours were immunoblotted with antibodies against the indicated proteins.

**Supplemental Figure S4.** Activation of M1C in patient-derived MGH1138 and MGH1112 cells. A and. **B.** MGH1138 cells treated with 50 nM adagrasib (**A**) and 50 nM RMC-7977 (**B**) for 48 hours were analyzed for M1C transcripts by qRT-PCR. The results (mean±SD of 4 determinations) are expressed as relative levels compared to that obtained for control cells (assigned a value of 1). **C and D.** GSEA of RNA-seq data from MGH1112 cells with M1C silencing using the indicated HALLMARK gene signatures. **E.** Lysates from MGH1112/tet-MUC1shRNA treated with DOX for 7 days were immunoblotted with antibodies against the indicated proteins.

**Supplemental Figure S5.** M1C ADC is effective against sotorasib-resistant cells. **A.** H358, HCC44 and H2122 cells treated with indicated concentrations of M1C ADC for 7 days were analyzed for cell viability by Alamar Blue staining. The results (mean±SD of six determinations) are expressed as relative cell number (% control) compared to that for untreated cells. Indicated are the M1C ADC IC50 values. **B.** H2122 cells treated with 30 nM M1C ADC for 7-14 days were analyzed for colony formation. Shown are representative photomicrographs of stained colonies (left). The results (mean±SD of three determinations) are expressed as relative colony formation compared to that for untreated cells (assigned a value of 1) (right). **C.** H2122 cells treated with vehicle or 50 nM M1C ADC for 7 days were analyzed for tumorsphere formation. Shown are representative photomicrographs of tumorspheres (left). The results (mean±SD of three determinations) are expressed as number of tumorsphere formations compared to that for vehicle treated cells (right). **D.** H358 cells left untreated or treated with 50 nM adagrasib for 2 days were analyzed by flow cytometry with a control IgG and anti-M1C. Shown are histograms of cells positive for M1C expression (left). The bar plot shows the mean fluorescence intensity (MFI), presented as the mean ± SD of three independent determinations (right). **E.** H358 cells treated with 50 nM adagrasib for 2 days and then M1C ADC for 7 days were analyzed for cell viability by Alamar Blue staining. The results (mean±SD of six determinations) are expressed as relative cell number (% control) compared to that for untreated cells. Indicated are the M1C ADC IC50 values. **F.** H2122 and H2122-SR cells treated with the indicated concentrations of M1C ADC for 7 days were analyzed for cell viability by Alamar Blue staining. The results (mean±SD of six determinations) are expressed as relative cell number (% control) compared to that for untreated cells. Indicated are the M1C ADC IC50 values. **G.** H2122-SR cells treated with 50 nM M1C ADC for 7 days were analyzed for colony formation. Shown are representative photomicrographs of stained colonies (left). The results (mean±SD of three determinations) are expressed as relative colony formation compared to that for untreated cells (assigned a value of 1) (right). **H.** H2122-SR cells treated with vehicle or 50 nM M1C ADC for 7 days were analyzed for tumorsphere formation. Shown are representative photomicrographs of tumorspheres (left). The results (mean±SD of three determinations) are expressed as number of tumorsphere formations compared to that for vehicle treated cells (right). **I.** Mean body weight changes of H358-SR tumor bearing mice treated with vehicle or M1C ADC. The results are expressed as the mean ratio of relative body weight for which the SEMs were <10%.

**Supplemental Figure S6.** Effectiveness of the M1C ADC against MGH1112 cells and analysis of MUC1 expression in NSCLC KRAS G12C tumors. **A.** MGH1112 cells treated with 100 nM M1C ADC for 7 days were analyzed for colony formation. Shown are representative photomicrographs of stained colonies (left). The results (mean±SD of three determinations) are expressed as relative colony formation compared to that for untreated cells (assigned a value of 1) (right). **B.** MGH1112 cells were treated with the indicated concentrations of M1C ADC and sotorasib for 7 days. Cell viability was assessed by Alamar Blue staining (mean of three determinations). Indicated are the combination indices determined using HSA scores. **C.** Mean body weight changes of MGH1112 tumor bearing mice treated with vehicle or M1C ADC. The results are expressed as the mean ratio of relative body weight for which the SEMs were <10%. **D.** H358/tet-MUC1shRNA cells treated with vehicle or DOX for 7 days were analyzed for the indicated transcripts. The results (mean±SD of 4 determinations) are expressed as relative levels compared to that obtained for vehicle-treated cells (assigned a value of 1**). E.** MGH1112/tet-MUC1shRNA cells treated with vehicle or DOX for 7 days were analyzed for the indicated transcripts. The results (mean±SD of 4 determinations) are expressed as relative levels compared to that obtained for vehicle-treated cells (assigned a value of 1**). F**. Correlation of mucin and TFF1 expression in the mucinous signature from the KRYSTAL-1 trial.

**Supplemental Table S1.**
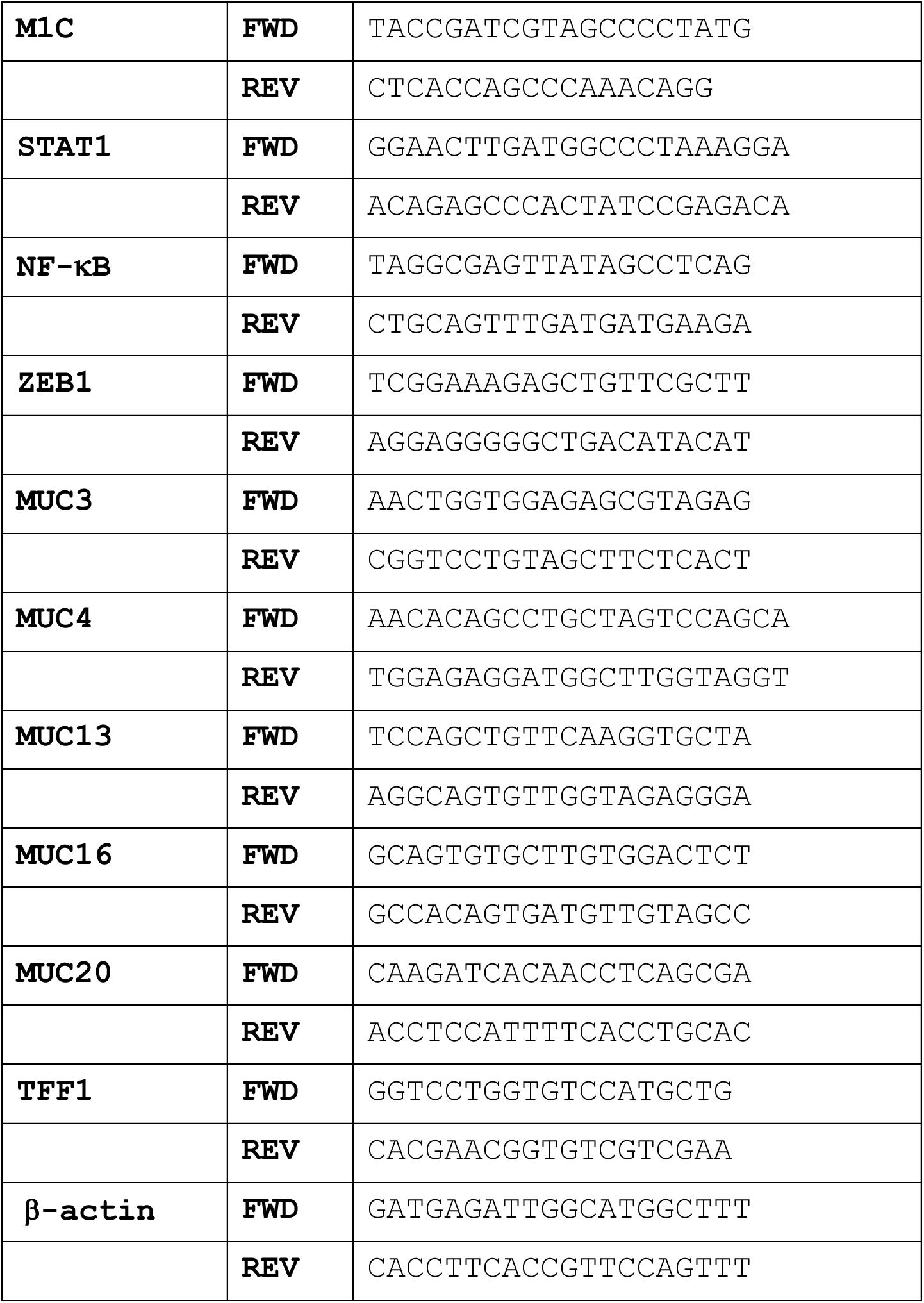
Primers used for qRT-PCR analysis.

## Notes

### Competing Interest Statement

The authors have declared no competing interest.

