## Supplementary Figures for "M1C is a druggable target for NSCLC KRAS G12C mutant tumors resistant to KRAS inhibitors"

### Supplementary Figure 1

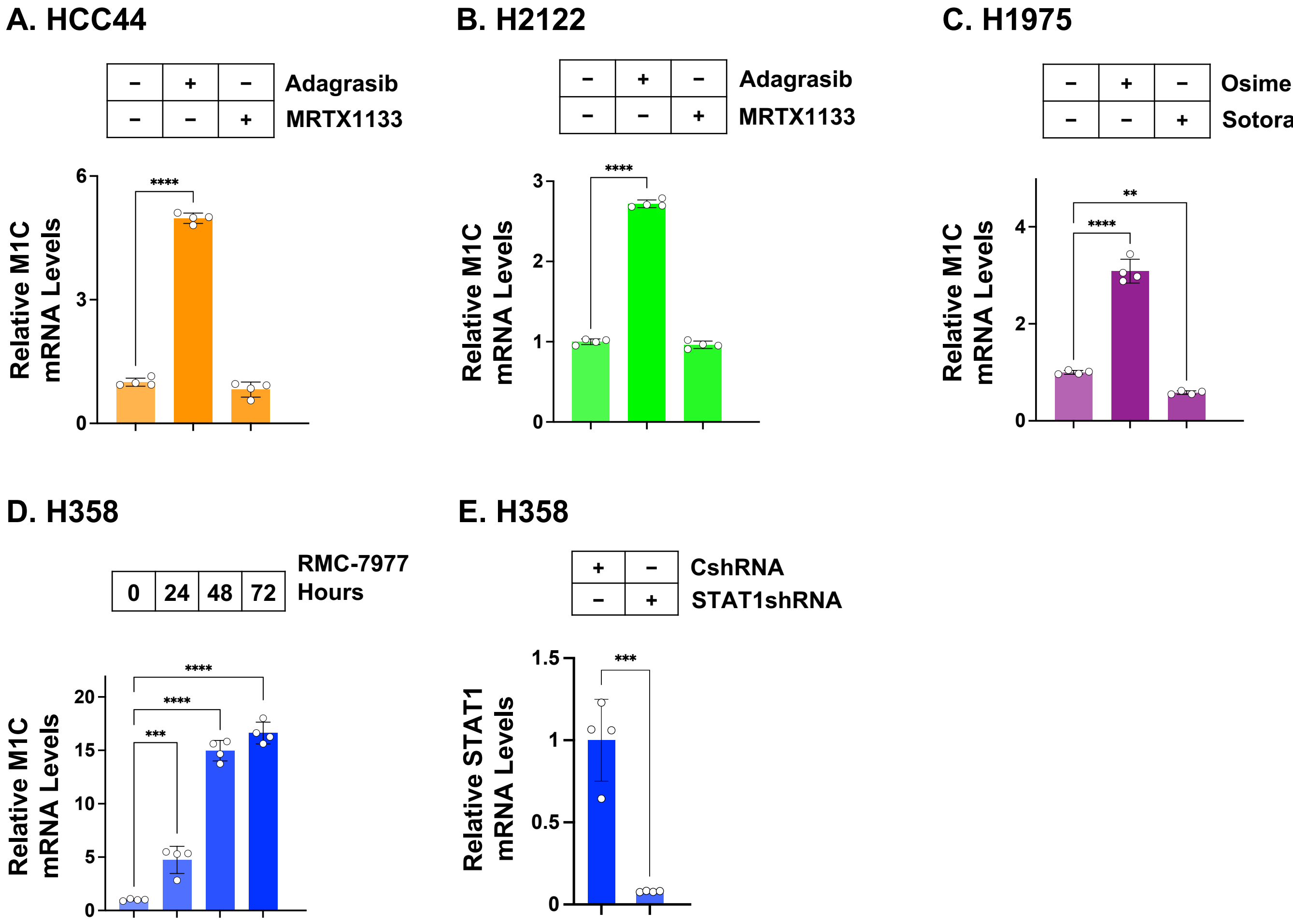

#### Supplementary Figure 2

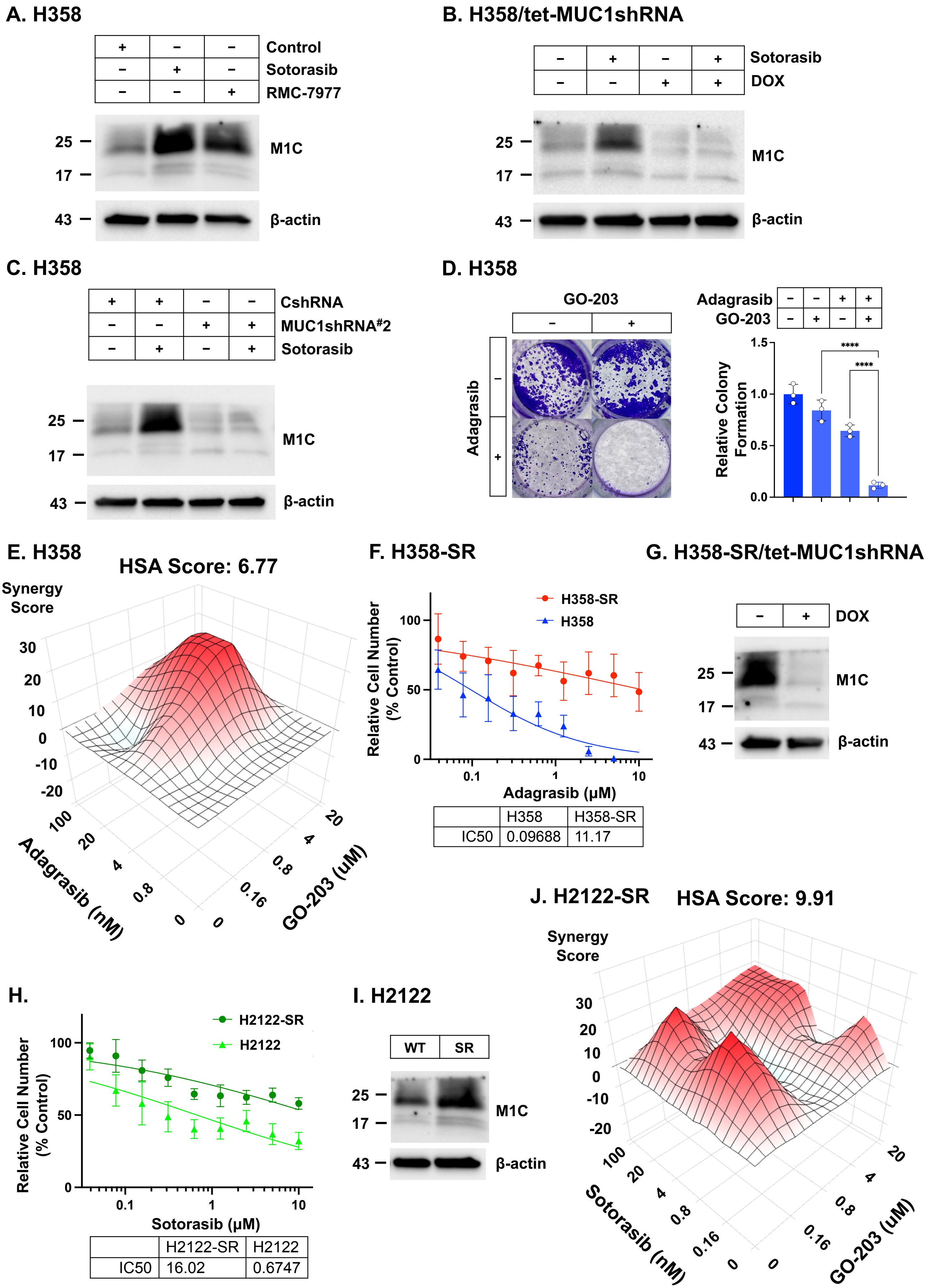


| | H358 | H358-SR |
|---|---|---|
| IC50 | 0.09688 | 11.17 |

| | H2122-SR | H2122 |
|---|---|---|
| IC50 | 16.02 | 0.6747 |

Supplementary Figure 3

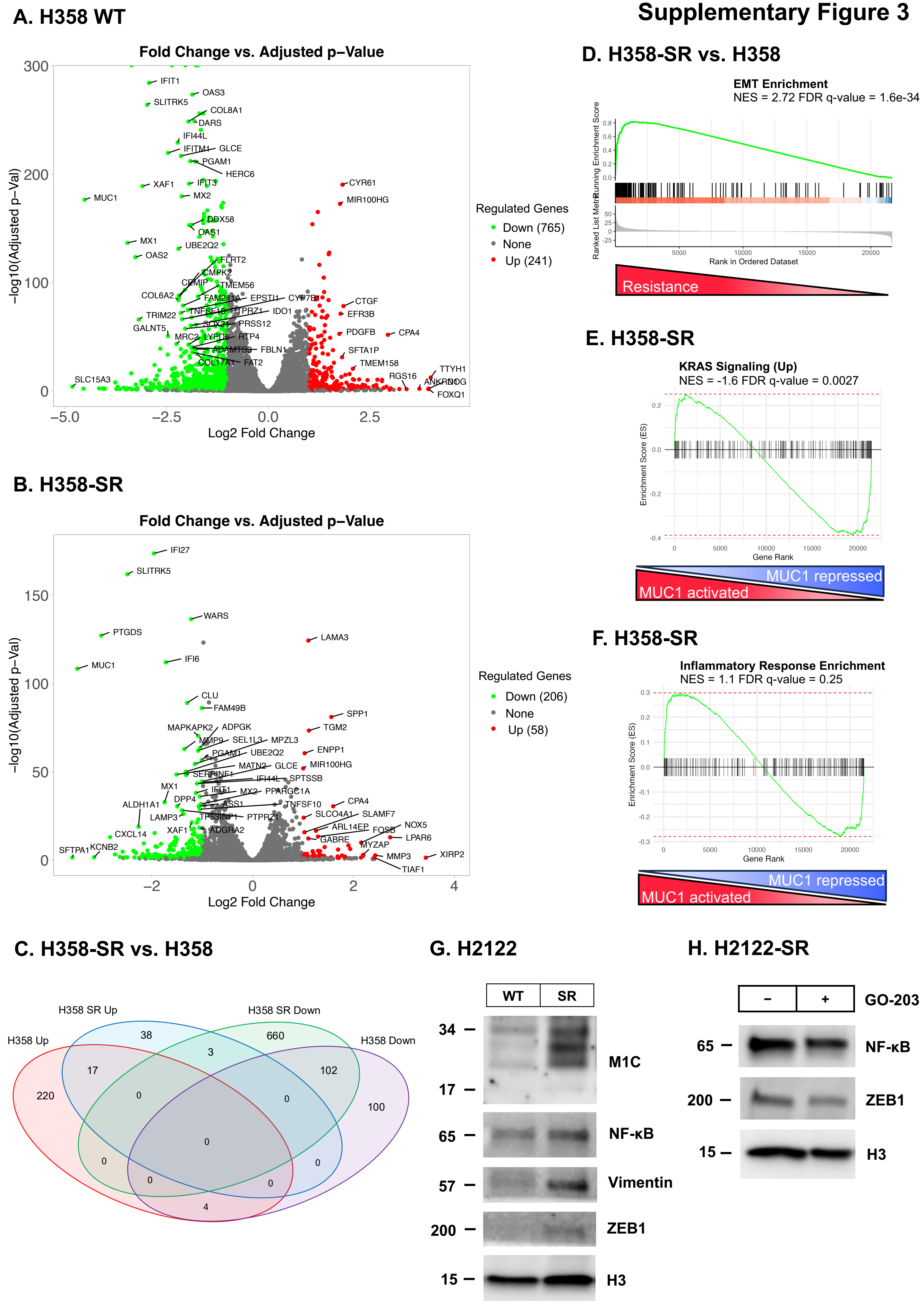

#### Supplementary Figure 4

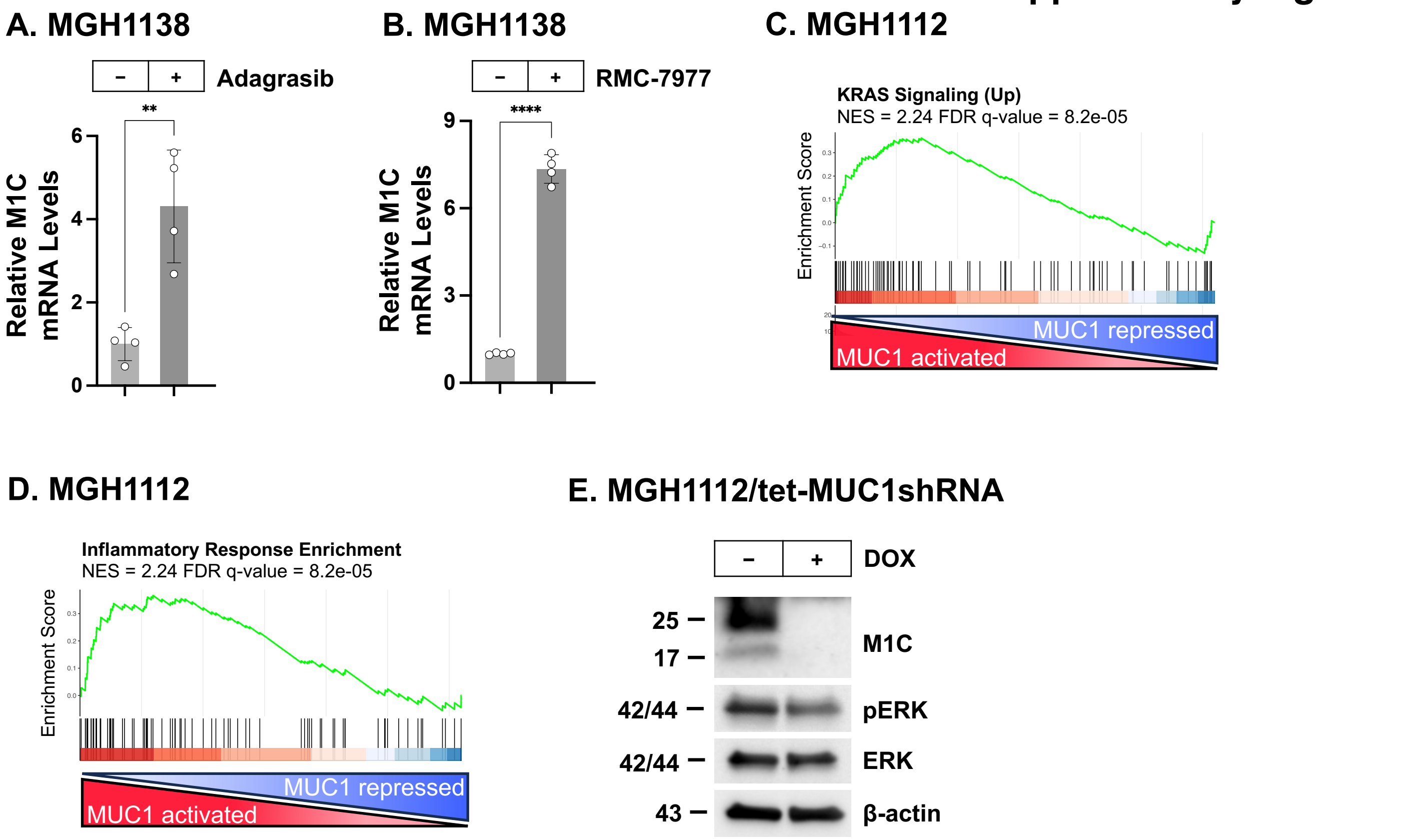

### Supplementary Figure 5

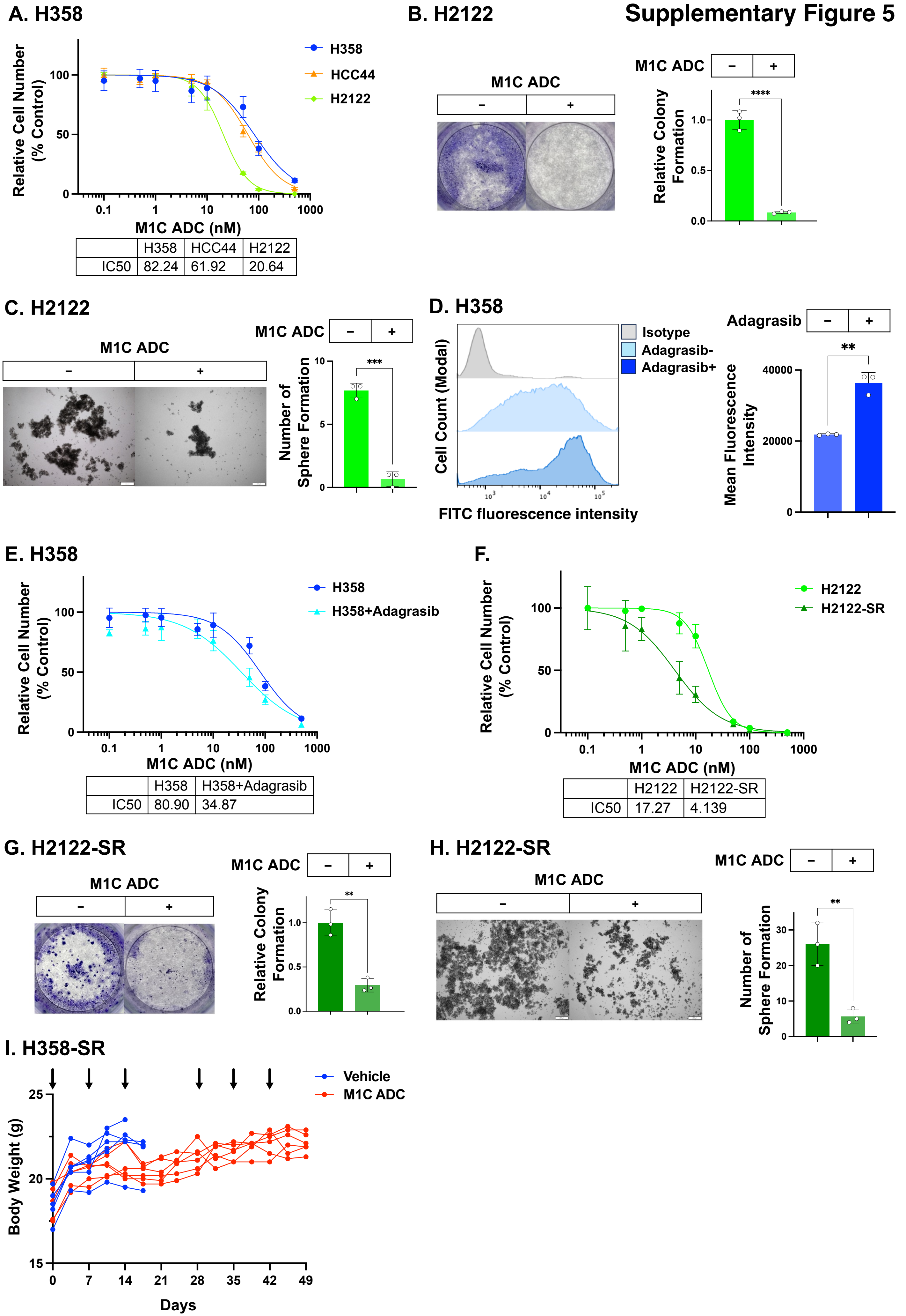

#### Supplementary Figure 6

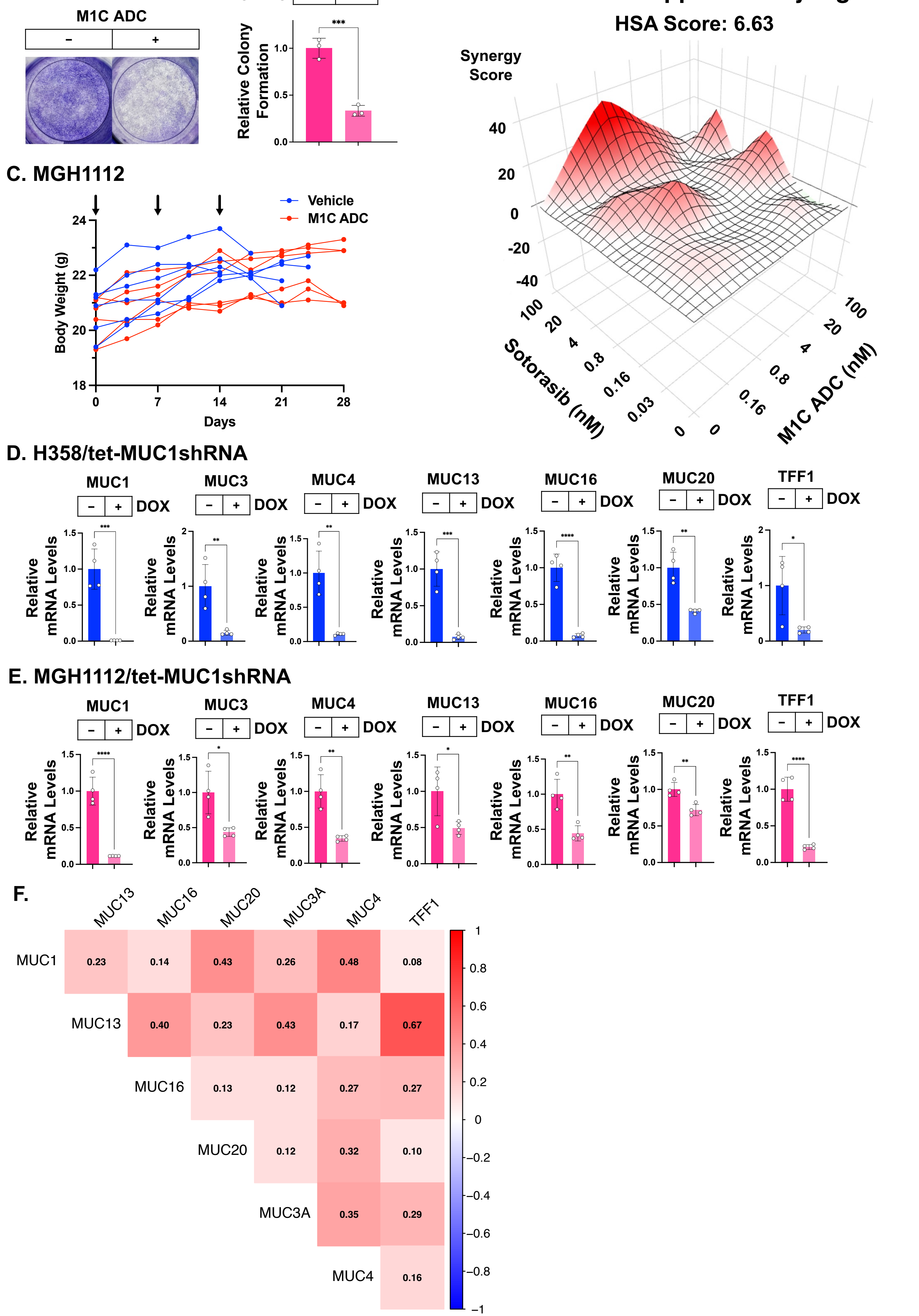
